# Improving Multichannel Raw Electroencephalography-based Diagnosis of Major Depressive Disorder via Transfer Learning with Single Channel Sleep Stage Data*

**DOI:** 10.1101/2023.04.29.538813

**Authors:** Charles A. Ellis, Abhinav Sattiraju, Robyn L. Miller, Vince D. Calhoun

## Abstract

As the field of deep learning has grown in recent years, its application to the domain of raw resting-state electroencephalography (EEG) has also increased. Relative to traditional machine learning methods or deep learning methods applied to manually engineered features, there are fewer methods for developing deep learning models on small raw EEG datasets. One potential approach for enhancing deep learning performance, in this case, is the use of transfer learning. While a number of studies have presented transfer learning approaches for manually engineered EEG features, relatively few approaches have been developed for raw resting-state EEG. In this study, we propose a novel EEG transfer learning approach wherein we first train a model on a large publicly available single-channel sleep stage classification dataset. We then use the learned representations to develop a classifier for automated major depressive disorder diagnosis with raw multichannel EEG. Statistical testing reveals that our approach significantly improves the performance of our model (p < 0.05), and we also find that the performance of our approach exceeds that of many previous studies using both engineered features and raw EEG. We further examine how transfer learning affected the representations learned by the model through a pair of explainability analyses, identifying key frequency bands and channels utilized across models. Our proposed approach represents a significant step forward for the domain of raw resting-state EEG classification and has broader implications for use with other electrophysiology and time-series modalities. Importantly, it has the potential to expand the use of deep learning methods across a greater variety of raw EEG datasets and lead to the development of more reliable EEG classifiers.

## I. Introduction

The application of deep learning methods to raw resting-state electroencephalogram (EEG) data has grown increasingly common in recent years. However, given the relatively small size of many locally collected EEG datasets and the expense of collecting larger datasets, it can be challenging to train reliable EEG deep learning models. It would be ideal if large existing EEG datasets could be leveraged to improve performance in other EEG deep learning applications. Other areas of deep learning (e.g., image classification) often use transfer learning for that reason. Additionally, some EEG studies involving manually extracted features (e.g., spectrograms) have developed transfer learning approaches [1], [2]. Nevertheless, while some initial efforts have been made to develop pretraining methods for EEG, there remains significant opportunities for growth in that arena. In this study, we implement a novel approach to demonstrate how large publicly available raw single-channel EEG sleep stage classification datasets can be used to pretrain a convolutional neural network (CNN) for automated multichannel EEG-based major depressive disorder (MDD) diagnosis. We show that the representations learned by the pretrained CNN can be used to obtain well-above chance-level performance without any further tuning of the convolutional layers, and we further show how our pretrained model can yield higher performance when fully tuned relative to the same architecture without pretraining. We lastly apply explainability analyses to examine how pretraining affected the learning of representations for canonical frequency bands and channels.

Historically, a large number of studies have applied machine learning and deep learning methods to extracted or manually engineered EEG features [3], [4]. While these approaches can be effective, their use of manually extracted feature inherently limits the size of the feature space from which they can learn. As such, the application of deep learning methods to raw EEG has become increasingly common, as deep learning methods like CNNs have the capability to perform automated feature extraction [5], [6].

Unfortunately, relative to traditional machine learning models trained on manually extracted features, deep learning models require larger datasets to be effective. Large datasets can be very time- and resource-intensive to collect and may be outside the capabilities of smaller research centers. Within the space of raw EEG analysis, some studies have addressed this problem via the use of data augmentation with additive noise and segmentation with large overlaps between samples [7][8]. Another possible solution is the use of transfer learning.

Transfer learning involves the training of a model on a large dataset from one task. The representations learned from that dataset can then be reused in the initialization of model weights for a second task. Transfer learning is common in applications like image classification [9], [10] and has been used in multiple studies involving extracted EEG features [1], [2]. Nevertheless, relatively few pretraining approaches for raw EEG have been developed. Of those that have been developed, they often involve event-related potentials [11] or task data [12] that are very different from resting state data or the use of self-supervised learning [13]. One study of particular interest used a pretrained architecture originally developed in the imaging domain to classify raw EEG data [14].

In this study, we develop an approach for pretraining models on publicly available raw one-channel sleep stage classification data and subsequently applying those models for the classification of multichannel EEG data. We demonstrate our approach within the context of automated MDD diagnosis. We find that our approach is able to learn generalizable features and that our pretraining approach significantly (p < 0.05) improves model performance relative to a baseline model with the same architecture and a traditional training approach. Lastly, we apply explainability approaches to examine differences in the spectral and spatial features learned by each of our models.

## II. Methods

In this study, we used two publicly available datasets: a sleep staging dataset and a MDD diagnosis dataset. We trained a model on the sleep stage dataset and used the resulting model weights to initialize several pretrained models. We compared the performance of our pretrained models to a baseline model via statistical testing and lastly applied spectral and spatial explainability approaches to gain insight into our models. Additionally, in an effort to ensure reproducibility of our results as well as enable other researchers to use our pretrained models, we have made our code and pretrained sleep models available on GitHub (https://github.com/cae67/Pretraining/).

### A. Datasets

We used two publicly available datasets in this study: a single-channel EEG sleep stage classification dataset and a multichannel EEG MDD dataset. Given that both datasets were publicly available, it was unnecessary to obtain approval for this study from an institutional review board.

#### Sleep Stage Dataset

We used a portion of the Sleep Cassette data from the Sleep-EDF Expanded dataset [15] on PhysioNet [16]. The dataset was composed of 153 20-hour recordings from 78 study participants, though due to computational constraints in the training process we used data from only 39 participants. The data was recorded at a sampling rate of 100 Hertz. While data from 2 electrodes were collected, we used data from the Fpz-Cz electrode. Thirty-second segments of data were assigned to Awake, Rapid Eye Movement (REM), Non-REM 1 (NREM1), NEM2, NREM3, and NREM4 sleep stages by an expert.

Based on the expert annotations, we separated the data into 30-second segments. To alleviate class imbalances, we removed the Awake data at the start of each recording and many Awake segments at the end of each recording. According to existing clinical practice [17], we reassigned NREM4 to NREM3. We lastly z-scored samples from each recording. Our final dataset was composed of 37,605, 7,534, 35,262, 7,939, and 13,801 Awake, NREM1, NREM2, NREM3, and REM samples, respectively.

#### MDD Dataset

We used a scalp EEG dataset [18] composed of 5-10 minute resting-state recordings with eyes closed from 28 healthy controls (HCs) and 30 individuals with MDD (MDDs). Recordings were performed at a sampling frequency of 256 Hertz in the standard 10-20 format with 64 electrodes.

Similar to previous MDD studies, we used 19 channels: Fp1, Fp2, F7, F3, Fz, F4, F8, T3, C3, Cz, C4, T4, T5, P3, Pz, P4, T6, O1, and O2. We performed several data harmonization steps to make the MDD data similar to the sleep data that would be used for pretraining. Specifically, we downsampled the data to 100 Hz and channel-wise z-scored each recording separately. To increase the size of the dataset available for training, we used a 30-second sliding window with a 2.5-second step size to epoch the data. Our final dataset had 2840 HCs and 2850 MDDs.

### B. Model Development

We adapted the architecture (shown in Fig. 1) that was used for MDD classification in [8], [19] and originally for schizophrenia classification in [5]. We trained 4 models using highly similar architectures: a sleep stage model (Model S) that would later be used for pretraining MDD models, a baseline MDD model with no pretraining (Model A), a pretrained MDD model with frozen feature extraction layers (Model B) to examine the transferability of the features learned by the sleep model, a pretrained model (Model C) that extended Model B by using the trained dense layer weights and training the whole network, and a pretrained MDD model with no frozen feature extraction layers (Model D). MDD models used sensitivity (SENS), specificity (SPEC), and balanced accuracy (BACC) when assessing model performance, and the Model S used SENS, precision (PREC), and the F1 score.

**Fig. 1.**
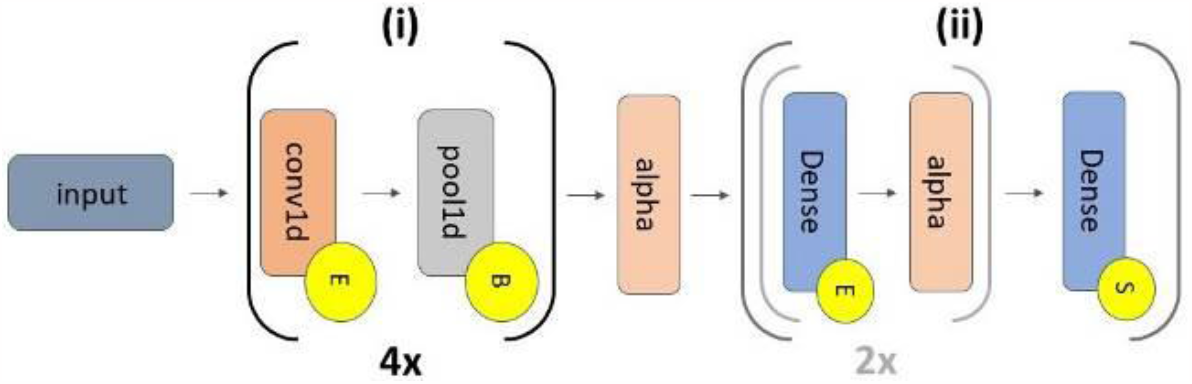
Model Architecture. Models A through D have 2 sections that are separated by an alpha dropout layer (alpha): (i) feature extraction, which repeats 4 times, and (ii) classification. The grey inset within section (ii) repeats twice. The 4 convolutional (conv1d) layers have 5, 10, 10, and 15 filters, respectively (kernel size = 10, 10, 10, 5). They are followed by max pooling layers (pool size = 2, stride = 2). Section (ii) has 3 dense layers (64, 32, and 2 nodes) with interspersed alpha dropout layers (alpha) with rates of 0.5. The alpha layer between (i) and (ii) also has a rate of 0.5. Yellow circles containing an “E”, “B”, or “S” correspond to ELU activations, batch normalization, and softmax activations. All conv1d and dense layers had max norm kernel constraints with a max value of 1. In Model S the channel dropout layer was placed between “Input” and Section (i).

#### 1) Model S

When developing the sleep model for pretraining, our approach involved several key components. We first created 19 duplicates of each 30-second sleep segment and added Gaussian noise (µ=0, σ=0.7) to each duplicate. After concatenating each duplicate, samples input to the model were of the same dimensions of those of the MDD models (i.e., 3000 time points by 19 channels). Our next key component was a channel dropout layer [20] with a dropout rate of 25% that we added at the beginning of the model to force the model to learn representations from all input channels rather than overfitting on only one channel. We used a training approach similar to that of [21]. We used a subject-wise group shuffle split cross validation with an 80-10-10 training-validation-test split. We trained the model for a maximum of 200 epochs with a batch size of 128, an Adam optimizer (learning rate = 0.0075) with an adaptive learning rate that decreased by an order of magnitude after every 5 epochs without an increase in validation accuracy, and early stopping after 20 epochs without an increase in validation accuracy.

#### 2) Model A

We used 25-fold subject-wise stratified group shuffle split cross-validation to prevent samples from the same participant being distributed across training, validation, and test sets. Training, validation, and test sets were assigned approximately 75%, 15%, and 10% of the data in each fold, respectively. To increase dataset size, we applied a data augmentation approach that involved creating a copy of the training data in each fold, adding Gaussian (µ=0, σ=0.7) noise to the copy, and using the combined original and augmented data for training. This approach has been used in multiple previous studies [7]. We next used a class-weighted loss function to account for training set class imbalances. We applied the Adam optimizer (learning rate = 0.0075), a batch size of 128, 35 epochs, and early stopping if 10 epochs passed without an increase in validation accuracy.

#### 3) Model B

To examine the utility of the features learned by Model S, we used the Model S weights from each fold, removed the channel dropout layer at the start of the model, froze the convolutional layer weights, re-initialized the dense layer weights, and decreased the number of nodes for the final softmax layer from 5 to 2. After making these modifications to the pre-trained model, we used a virtually identical training approach to that used in the training of Model A. However, in contrast to the Model A training, we repeated the training once per sleep fold (i.e., 10 times) resulting in 250 models (i.e., 10 sleep folds x 25 MDD folds).

#### 4) Model C

After examining whether Model B had useful representations, we further optimized Model B by unfreezing all weights to see if performance could be further improved. We initialized Model C with the Model B weights and otherwise used the same training procedure as in Models A and B. This yielded 250 models (i.e., 10 sleep folds x 25 MDD folds).

#### 5) Model D

To examine whether the staged training process of Models B and C was better relative to just training directly with the Model S weights, we used the Model S weights from each fold, removed the channel dropout layer at the start of the model, re-initialized the dense layer weights, and decreased the number of nodes for the final softmax layer from 5 to 2. After making these modifications to the pre-trained model, we used an identical training approach to that used in the training of Model B. This yielded 250 models (i.e., 10 sleep folds x 25 MDD folds).

### C. Model Performance Analysis

After training each of the models, we wanted to determine whether pretraining improved baseline model performance in a statistically significant manner. To that end, we generated 10 duplicates of the Model A performance (i.e., one per sleep fold) and performed family-wise, paired, two-tailed t-tests between the BACC, SENS, and SPEC of each MDD model. Afterwards, we performed false discovery rate correction [22] of all p-values (α = 0.05) and identified significant tests (p < 0.05).

### D. Explainability Analysis

We lastly examined how pretraining affected the representations learned by MDD models. Specifically, we applied an adaptation of a spectral explainability approach that was presented in previous works [23], [24] and a zero-out channel ablation approach [25], [26] to the MDD models.

Our spectral explainability approach involved converting test samples to the frequency domain with a fast Fourier transform (FFT), replacing the FFT coefficients within a given frequency band with zeros, and converting back to the time domain, and calculating the percent change in BACC following perturbation. We used the δ (0-4 Hz), θ (4-8 Hz), α (8-11 Hz), β (12-25 Hz), and γ (25-50 Hz) frequency bands. It would ordinarily be best to permute frequency coefficients multiple times across samples within a given band. However, due to the relatively large number of models that we analyzed (i.e., 775 models), doing so was computationally prohibitive, and we opted for simply zeroing out individual frequency bands.

Our channel ablation approach involved iteratively assigning each channel in test samples to values of zero and examining the change in model performance. While we would have preferred to use the line-noise ablation approach described in [26], which can amplify the effects of ablation upon the model, the use of publicly available data with a notch filter at 50 Hz removed line-related noise. As such, the model would not learn to consider line-related noise to be neutral information, and the use of line-noise ablation would no longer presented significant advantages.

## III. Results and Discussion

In this section, we describe and discuss our model performance and explainability results within the context of existing literature. We further discuss some of the key aspects and broader implications of our approach, and lastly summarize the limitations and future work related to this study.

### A. Model Performance Comparison

Tables 1 and 2 show our Model S and MDD model performance results, respectively. Additionally, Fig. 2 shows the results for our t-tests comparing MDD model performance. Model S obtained modest levels of performance below that of existing sleep stage classification studies [21], [24]. Nevertheless, classification performance was above chance-level for most classes. This is important as it shows that our use of duplicate channels with additive noise and channel dropout did not prevent the model from learning.

**Table 1.**
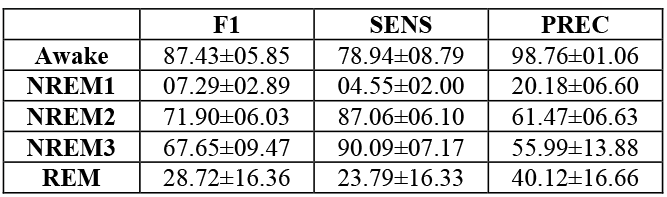
Model S Performance Results.

|  | F1 | SENS | PREC |
| --- | --- | --- | --- |
| **Awake** | 87.43±05.85 | 78.94±08.79 | 98.76±01.06 |
| **NREM1** | 07.29±02.89 | 04.55±02.00 | 20.18±06.60 |
| **NREM2** | 71.90±06.03 | 87.06±06.10 | 61.47±06.63 |
| **NREM3** | 67.65±09.47 | 90.09±07.17 | 55.99±13.88 |
| **REM** | 28.72±16.36 | 23.79±16.33 | 40.12±16.66 |

**Table 2.** Performance Results for MDD Models.

|  | BACC | SENS | SPEC |
| --- | --- | --- | --- |
| **Model A** | 83.57±15.13 | 91.11±14.44 | 76.03±31.72 |
| **Model B** | 86.00±13.79 | 88.63±16.29 | 83.36±24.74 |
| **Model C** | 87.16±13.40 | 91.01±12.98 | 83.30±25.71 |
| **Model D** | 85.68±13.89 | 88.25±16.27 | 83.11±24.89 |

**Fig. 2.**
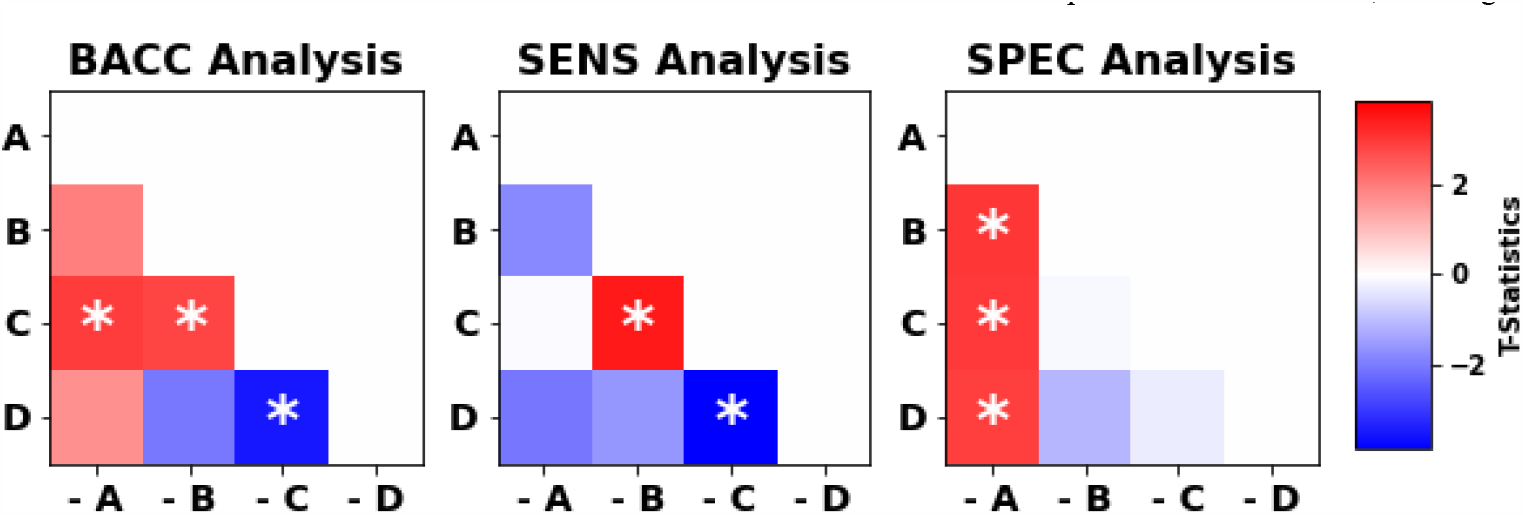
Model Performance Test Analysis. The panels show differences in BACC, SENS, and SPEC between models, respectively. The y-axis indicates a given model, and the x-axis indicates corresponding models that are subtracted from the models on the x-axis in the t-tests. T-statistics for each test are displayed in each panel, and the t-statistics in each panel share the colorbar to the right of the figure. White asterisks indicate t-tests with statistically significant results following FDR correction (p < 0.05). Note that Model C obtains highest BACC, Model C has higher SENS than Models B and D, and all pretrained models (i.e., Models B through D) have higher SPEC than Model A.

Model A shows the baseline level of performance that could be expected from our architecture and data (Table 2). All metrics were above 75%, with SENS being significantly higher than SPEC and BACC being close to 84%. Fig. 2 shows the results of our statistical testing comparing the BACC, SENS, and SPEC of each of the models. Interestingly, model B, which just used the features extracted by model B, was able to obtain a BACC greater than, though not significantly greater than, model A. This indicates that the feature representations learned by Model S from the sleep data were rich enough to generalize to other domains. Model C was our highest performing model, obtaining significantly higher BACC (p < 0.05) than all of the other MDD models. It had comparable SPEC to Models B and D (p > 0.05) and SENS to model A. Moreover, it had significantly greater SENS than Models C and D (p < 0.05). Relative to the Model A baseline, pretraining improved model performance by more than 3.5% on average in a task for which model performance was already high. Our final model, Model D, demonstrated a nonsignificant increase in BACC relative to Model A. Of note, all pretrained MDD models (i.e., Models B through D) obtained greater SPEC than Model A (p < 0.05), and while the baseline Model A obtained greater SENS than Models B and D, that difference was nonsignificant (p > 0.05).

Other classification studies have used the same MDD dataset, so it is also important to assess how our pretraining performance compares to the performance reported in those studies. Relative to [8], our model demonstrated a marked improvement in performance across metrics, and relative to [19], our approach demonstrated a modest increase in performance. Both of these studies used raw EEG data. It should be noted that these studies are not totally comparable, as we used a much larger number of folds and models (i.e., 10 MDD folds resulting in 10 MDD models versus 10 sleep folds x 25 MDD folds, resulting in 250 MDD models). Additionally, relative to [27], a study that used a robust cross-validation approach and manually engineered features, we also demonstrated an improvement. A number of studies using raw EEG [28], [29] and manually engineered features [30]–[33] did obtain higher classification performance than our approach. However, their cross-validation approaches are either not clearly described, or it seems that they employed cross-validation approaches that allowed data leakage between the training and test sets. Namely, they seemed to allow samples from the same subjects, which had high levels of dependency, to be placed in both training and test sets within the same folds, which likely overinflated their model performance. This is unfortunately a common problem within the field and has been discussed in previous studies [27]. Our subject-wise, grouped cross-validation approach prevented this mixture of samples across training and test sets. As such, relative to studies using robust cross-validation approaches, our approach obtains comparable or higher levels of importance.

To complete our discussion of model performance, our comparison with previous studies in conjunction with our findings of improved performance in the results presented in this study suggest that pretraining deep learning models on single-channel sleep data is able to yield useful representations that are generalizable to other domains and that our pretraining approach can significantly improve model performance.

### B. Explainability Analysis

As shown in Fig. 3, all models greatly relied upon δ, β, and γ activity and generally ignored θ and α activity. The findings of δ, β, and γ importance fit with existing literature [8], [19]. However, previous explainability analyses (using different approaches) [8], [19] also identified α importance for MDD diagnosis, which contradicts our findings. Interestingly, while Model A tended to be more sensitive to β and γ perturbation than δ perturbation, Models B through D tended to be more sensitive to δ perturbation than to β and γ perturbation. Given how these changes in importance also coincided with increased MDD model performance, this could indicate that Model S was able to learn representations for δ activity from the sleep data that the MDD models were able to capitalize on. The ability of Model S to learn δ activity would fit with existing domain knowledge, as the importance of δ activity for sleep stage classification has been demonstrated in a variety of previous studies [21], [24], [34]–[36]. Moreover, δ is used clinically to identify the NREM3 sleep stage [17], for which Model S obtained a high level of performance. Additionally, given the relatively long length of δ waveforms compared to higher frequency β and γ waveforms, it makes sense that it would be easier for Model A to learn shorter, higher frequency waveforms and thus quickly overfit on that activity while missing out on the lower frequency activity that Model S provided Models B through D.

**Fig. 3.**
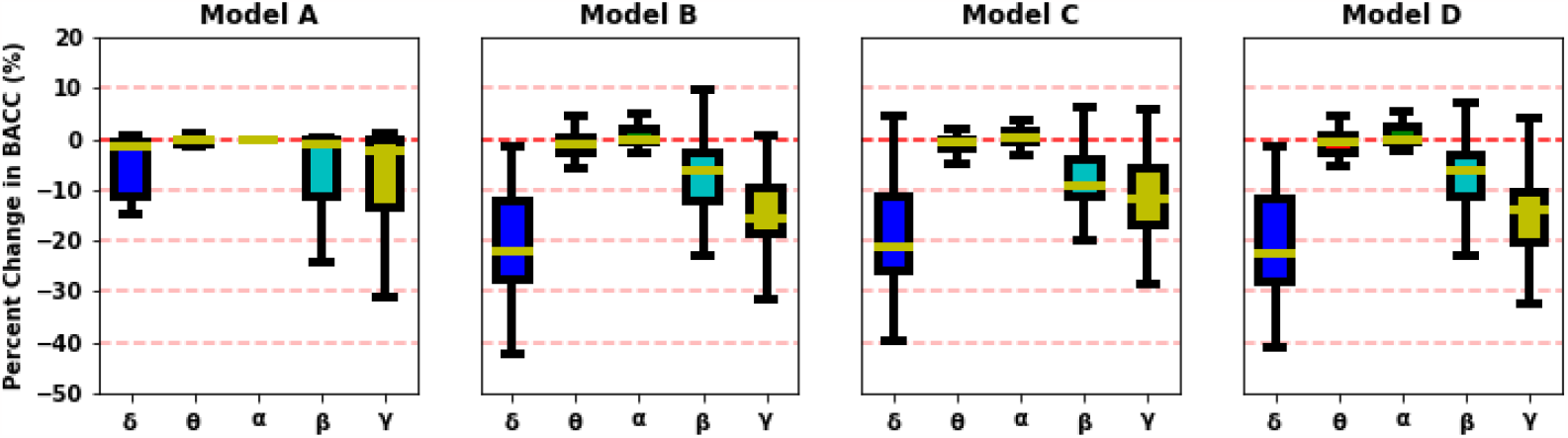
Spectral Importance. In panels from left to right are spectral importance for Models A, C, and D. The presence of 25 Model A models versus 250 models for Models B through D resulted in large differences in variance of model importance. To have comparable levels of variance in our display, we calculated the mean change in BACC within each set of 10 sleep folds for each of the 25 MDD folds in Models B through D. The y-axes are aligned and show the percent change in BACC, and the x-axes indicate each frequency band. Note that δ, followed by γ activity was important across models, though the pretrained models also prioritized β activity.

There were also some similarities in spatial importance, as shown in Fig. 4. Namely, F7, Pz, and C3 were of importance across models. Additionally, the pretrained models seemed to generally prioritize channels in manner similar to one another. Nevertheless, there were some key differences between Model A and the pretrained models. The pretrained models generally relied most upon Fz, while Model A did not rely upon Fz. In contrast to the other models, Model C also greatly prioritized P3. Additionally, while the removal of many channels improved the pretrained model performance, removal of channels from Model A generally decreased performance. Models B through D were also much more sensitive to channel loss than Model A and had a much wider range in the effect of perturbation. Model C, in particular, seemed to be more sensitive to channel loss than Models B and D. This key reliance upon frontal and parietal electrodes is similar to findings of existing studies [8], [19].

**Fig. 4.**
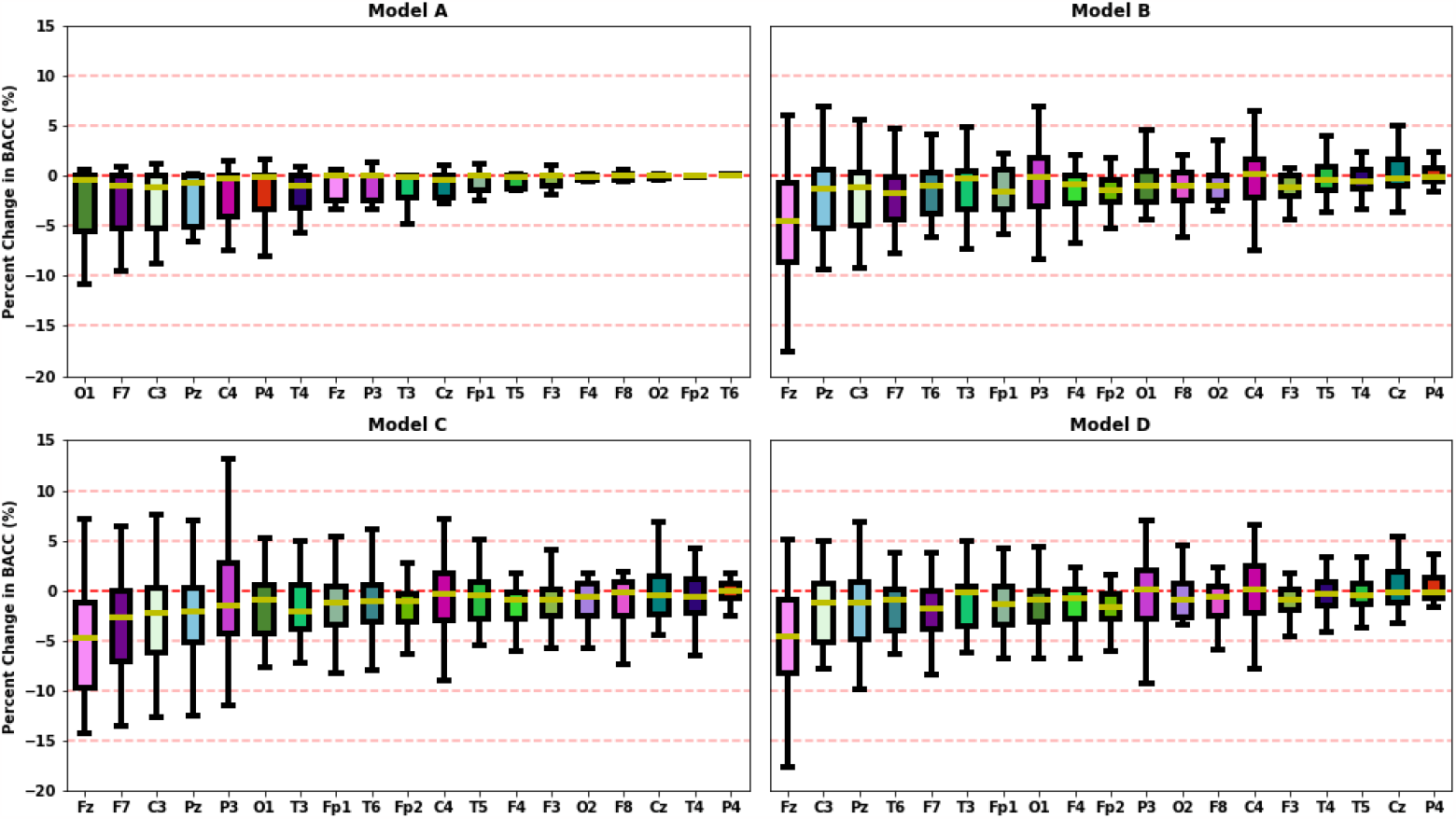
Spatial Importance. The top left, top right, bottom left, and bottom right panels show spatial importance for Models A, B, C, and D, respectively. The presence of 25 Model A models versus 250 models for Models B through D resulted in large differences in variance of model importance. To have comparable levels of variance in our display, we calculated the mean change in BACC within each set of 10 sleep folds for each of the 25 MDD folds in Models B through D. Channels are displayed on the x-axis, and the percent change in BACC is shown on the y-axis. Due to the relatively low values for Model A, the channels in each panel displayed in order of increasing 25% quantiles from left to right. Note how Pz, F7, and C3 are generally important across models. Additionally, the pretraining models, Models B through D, relied greatly upon Fz. It should also be noted how there were relative few Model A models for which removing a given channel improved model performance. However, performance for Models B through D tended to vary more in response to channel loss. Model C, the highest performing model tended to vary most in response to channel loss.

### C. Further Discussion on Approach Development

Our pretraining approach had several key aspects that enabled it to improve automated MDD diagnosis performance.

1. We used sleep stage data in our pretraining approach. A number of large sleep stage datasets are publicly available, which enhances the potential for widespread adoption and impact of our approach [15], [37], [38]. While we could have used smaller, multichannel EEG datasets that corresponded to other classification tasks (e.g., diagnosis of schizophrenia [7] or Alzheimer’s disease [39]), we assumed that sleep stage data would be more useful for model pretraining. Whereas other EEG tasks often have specific frequency bands of importance [7], virtually all EEG frequency bands are used clinically for identifying sleep stages [17] and have been identified as relevant in previous EEG sleep stage classification studies involving explainability methods [21], [24], [34]–[36]. If we were to use data from another EEG task for pretraining, there is a strong possibility that the model would overfit on some frequency bands and learn poor representations for others. Moreover, other EEG tasks often tend to focus on specific electrodes and brain regions, which would result in them learning poor representations for some channels. By using single-channel sleep data, we ensured that the model had cause to learn from all channels. This leads to the next unique aspect of our approach.
2. Namely, we replicated each of the sleep data samples to have a number of channels equivalent to that of the MDD data. This process does significantly increase the size of the sleep dataset and can be prohibitive depending upon access to computational resources. Nevertheless, there are approaches to efficiently implement this in one’s code to make it more viable. In our case, we only used half of the sleep dataset, and future iterations of the approach with more efficient coding might further enhance the effects of pretraining.
3. We augmented the replicated channels with Gaussian noise in an attempt to more accurately model how EEG data might be expected to vary across channels and to force the sleep model to learn generalizable representations. Future studies might examine how different data augmentation approaches might be used in this context.
4. We applied channel dropout to force the sleep model to learn useful representations across channels. In contrast to the data augmentation-based channel dropout approach presented in [40] which would increase the sleep dataset size to unmanageable levels, we used the layer-based channel dropout approach presented in [20]. Because this regularization approach prevents the model from quickly converging and overfitting on specific channels, it was also necessary to train the model for a large number of epochs. It should be noted that while we had a total possible number of 200 training epochs, we also ended training if performance did not improve following 20 epochs. This cutoff could be further optimized in future studies.

### C. Broader Implications of Approach

Our approach has the potential to significantly improve the robustness and generalizability of models trained on small datasets. By learning a diverse set of EEG features from sleep data, the likelihood of overfitting on noisy features in small datasets can be reduced. Given the financial and logistical difficulties of collecting large sets of patient data, this could be particularly advantageous both in research and clinical contexts. While we applied our approach within the context of automated MDD diagnosis, our approach could also easily be applied to other EEG applications. Moreover, given the variety of waveforms found in EEG sleep data, our approach has the potential to be applied within the context of other electrophysiology modalities like magnetoencephalography [41], electromyography [42], electrooculography [43], or multimodal sleep staging [25], [44]. This form of cross-modality pretraining could greatly accelerate model development across a variety of electrophysiology modalities. Additionally, while we believe the development of our approach represents a key early step for the development of pretraining approaches for EEG, we hope that it will inspire other researchers to develop creative EEG pretraining solutions. This could eventually culminate in out-of-the-box models like those frequently used in image classification [9], [10], and this is the primary motive for our release of our pretrained sleep models on GitHub (see link in Methods section). Within the context of broader time-series classification outside of electrophysiology, the general concept of using duplicated single-series data with rich representations could be applied to improve model performance in multi-time-series applications [45], [46].

### D. Summary of Limitations and Next Steps

While our pretraining approach increased model performance and we examined the effect of pretraining upon learned representations, there are several avenues for further development and investigation of our approach.

In future studies, we will examine how the performance of our model upon the initial sleep stage classification task affects subsequent MDD model performance. We will also examine how each aspect of our proposed approach contributes to final MDD model performance. For example, it is unclear which of the convolutional layers should be frozen when first training the pretrained models. In Model C, we trained on Model B weights for which all of the convolutional layer weights had originally been frozen, and in Model A we did not freeze any convolutional layer weights. Given the statistically significant differences in performance that we obtained for each of these approaches, it would seem that being forced to retain both high- and low-level features from different convolutional layers improved model performance. Nevertheless, systematically evaluating what levels of features should be retained could be helpful.

Additionally, our pretrained models seemed to be highly sensitive to channel loss. In future studies, investigating the utility of channel dropout for training using the pretrained features could potentially alleviate this sensitivity [20]. Along with this investigation of sensitivity to channel loss, recent studies have begun to investigate interactions learned between channels by deep learning models trained on EEG [8], [19]. Given the similarity of the replicated sleep channels that we train on in our approach, it is unclear how our approach might affect the interactions learned by the model between channels. Future studies might find this helpful to investigate. Lastly, the effects of MDD upon brain dynamics are spatially and spectrally widespread [27], which could affect the utility of our pretraining approach for MDD specifically. It would be instructive to apply our approach to other tasks like schizophrenia diagnosis wherein the effects of the disorder upon brain dynamics is more spatially localized [3], [7].

## IV. Conclusion

Deep learning methods are increasingly being applied to raw EEG data. However, the relatively small size of many datasets and the time and financial resources required to collect larger datasets make training robust and generalizable deep learning models a key technical challenge. In this study, we propose a new raw EEG transfer learning approach in which we transfer representations learned from single-channel sleep stage classification data to multichannel MDD diagnosis. Our approach improves model performance in a statistically significant manner (p < 0.05), and we show through explainability analyses how pretraining affects the features learned by our models. Importantly, our proposed approach has the potential to expand the use of deep learning methods across more raw EEG datasets and lead to the development of more reliable EEG classifiers. Moreover, it has broader implications for the acceleration of model development in other electrophysiology modalities. Our study represents a key step forward for the domain of raw EEG classification, making viable the application of deep learning methods to small EEG datasets. Moreover, we hope that it will galvanize the development and use of pretraining in raw EEG in future studies.

